# Integrated Spatial Transcriptomic and Proteomic Analysis of Fresh Frozen Tissue Based on Stereo-seq

**DOI:** 10.1101/2023.04.28.538364

**Authors:** Sha Liao, Yang Heng, Weiqing Liu, Jinqiong Xiang, Yong Ma, Ligen Chen, Xiuwen Feng, Dongmei Jia, Diyan Liang, Caili Huang, Jiajun Zhang, Min Jian, Kui Su, Mei Li, Yuin-Han Loh, Ao Chen, Xun Xu

**Affiliations:** BGI-Shenzhen, Shenzhen, Guangdong, China; BGI Research-Southwest, BGI, Chongqing 401329, China; JFL-BGI STOmics Center, Jinfeng Laboratory, Chongqing 401329, China; BGI Research Asia-Pacific, BGI, Singapore 138567, Singapore; Cell Fate Engineering and Therapeutics Lab, Cell Biology and Therapies Division, Institute of Molecular and Cell Biology (IMCB), Agency for Science, Technology and Research (A*STAR), 61 Biopolis Drive, Proteos, Singapore 138673, Republic of Singapore; Department of Biological Sciences, National University of Singapore, Singapore 117543, Singapore; NUS Graduate School’s Integrative Sciences and Engineering Programme, National University of Singapore, 28 Medical Drive, Singapore 117456, Singapore; Department of Physiology, Yong Loo Lin School of Medicine, National University of Singapore, Singapore 117593, Singapore

## Abstract

To simultaneously detect whole transcriptomes and protein markers on the same tissue section, we combined Cellular Indexing of Transcriptomes and Epitopes by Sequencing (CITE-seq) and Stereo-seq to develop the Stereo-CITE-seq workflow. Here, we demonstrated that Stereo-CITE-seq can co-detect mRNAs and proteins in immune organs with high spatial resolution, reproducibility and accuracy.

---

Cells within multicellular organism are spatially positioned, tightly regulated and highly connected with each other to maintain homeostasis in different tissue and organ systems. Taking advantage of single-cell RNA-seq technologies, cellular heterogeneity within tissues has been revealed at unprecedented single cell resolution ^1-4^. An inherent limitation of profiling tissues using single-cell RNA-seq is the lost of spatial information, which could not be captured during the sample processing steps. To this end, spatial RNA-seq has been developed to enable the comprehensive understanding of cellular and tissue physiology, within precise anatomical locations. Various techniques have been developed to provide spatial whole transcriptome analysis, including Visium ^5^, HDST ^6^ and Slide-seq ^7,8^. Nevertheless, these methods are confined by their lack of high resolution, and the failure to generate spatially resolved single-cell data. Hencetoforth, we have developed Stereo-seq, and successfully applied the technology for the transcriptional profiling of mouse embryos ^9^, zebra fish ^10^, axolotl brain ^11^, Arabidopsis leaves ^12^, at high spatial resolution (500nm), as compared to previous methods.

Studies have reported that mRNA abundance is not directly correlated with the protein abundance ^13,14^. Moreover, transcriptomic information alone could not faithfully delineate the functional microenvironments, which are essential in many diseases such as cancer ^15^. Co-detection methods such as Spatial PrOtein and Transcriptome Sequencing (SPOTS) ^16^, Spatial Multi-Omics (SM-Omics) ^17^, and Deterministic barcoding in tissue for spatial omics sequencing (DBiT-seq) ^18,19^ are currently used to integrate whole transcriptome and protein markers profiling on the same tissue section. On top of that, DBiT-seq has been further enhanced to enable the profiling of other omics layers, such as spatial ATAC-seq ^20^ and CUT&Tag ^21^. Unfortunately, the applicability of these methods are highly restricted by their low spatial resolution - 100 µm spot center-to-center distance for SPOTS, 200 µm for SM-Omics, 20 µm for DBIT-seq. Imaging-based spatial profiling methods such as co-detection by indexing (CODEX) ^22^, imaging mass cytometry (IMC) ^23,24^, Xenium ^25^ and CosMx^™^ Spatial Molecular Imager (SMI) ^26^ achieved high subcellular resolution, and are capable of concurrent detection of proteins and RNAs on the same tissue section ^27^. However, these techniques are limited by the lack of RNA detection multiplexity, and do not allow for whole transcriptome analysis. Thus, multi-omic profiling strategy with high spatial resolution and in-depth detection multiplexity is a gap that needs to be bridge, in order for the comprehensive understanding of biological processes in health and disease.

We integrated the Cellular Indexing of Transcriptomes and Epitopes by Sequencing (CITE-seq) and Stereo-seq workflows, and established the Stereo-CITE-seq technology, which enables simultaneously transcriptomic and proteomic profiling of tissue sample at high spatial resolution (Figure 1a, Methods). Similar to CITE-seq, Stereo-CITE-seq also relies on poly(dT) oligos on the Stereo-seq chip to capture antibody derived tag (ADT) and mRNA (Figure 1b, c). ADT and mRNA are then reverse-transcribed to obtain spatial information. After releasing, ADT products and cDNA will be separated by beads purification. Through PCR and sequencing, ADT and gene expression profiles are generated (Figure 1a).

**Figure 1.**
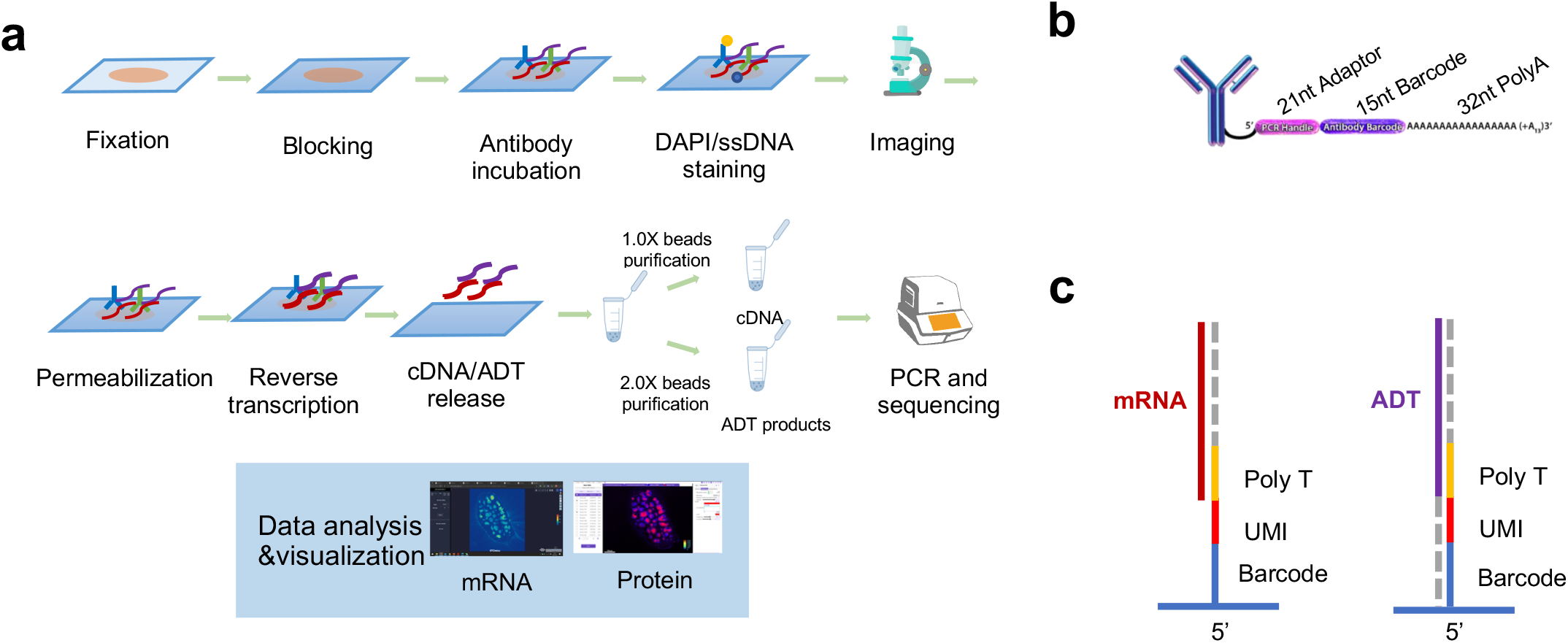
Principle of Stereo-CITE-seq. **a**, Overview of Stereo-CITE-seq workflow. **b**, Structure of antibody-derived tag (ADT). **c**, mRNA and ADT are both captured by Poly(dT) on the chip.

We first performed proteomics profiling on frozen mouse spleen section using 10 different antibodies for T-cells (CD3, CD4, CD8a), B-cells (CD19, CD45R, IgD), macrophages (CD169, CD163, F4/80) and fibroblasts (CD29). We also included two isotype control antibodies to detect non-specific background noise. In the mouse spleen sample, with the PCR duplication rate of 57.5%, the average protein UMI count per Bin 100 (50 × 50 µm area) is 16.1K, significantly higher than the ∼1024 average UMI count per spot (55 µm in diameter) reported for SPOTs ^16^ (Figure 2a). The average protein UMI count per Bin 50 (25 × 25 µm spatial area) is 4.05 K (Figure 2a), also higher than reported 885 average UMI count per 25 × 25 µm region reported in DBiT-seq ^18^. Next, we examined each ADT expression patterns without any additional filtering. The ADT expression patterns could nicely recapitulate the expected T cells, B cells, macrophages and fibroblasts spatial distribution and with insignificant background noise (Figure 2b). Notably, different ADT expression patterns were observed for CD4, CD8a, CD169 and F4/80, indicative that Stereo-CITE-seq could spatially resolve T cell and macrophage subpopulations (Figure 2b). Further, we assigned CD4, IgD, CD169 and CD163 ADT expression patterns with different colors and visualize their spatial location in the same image at Bin100 and Bin10 level (Figure 2c). We observed near-to-single cell level of spatial expressing patterns of T cells, B cells and macrophages at Bin 10 resolution, while considerable information was lost at Bin 100 level (Figure 2c), indicating spatial resolution is crucial for proteomic profiling. Non-supervised clustering on Bin10 spatial barcodes resulted in 6 clusters representing T cells (cluster1), B cells (Cluster 3) and macrophage subtype 1 (Cluster 2), macrophage subtype 2 (Cluster 4) (Figure 2d, 2e and 2f). Macrophage subsets (Cluster 2 and 4) could also be identified through non-supervised clustering, indicating the robustness of the ADT detection (Figure 2f).

**Figure 2.**
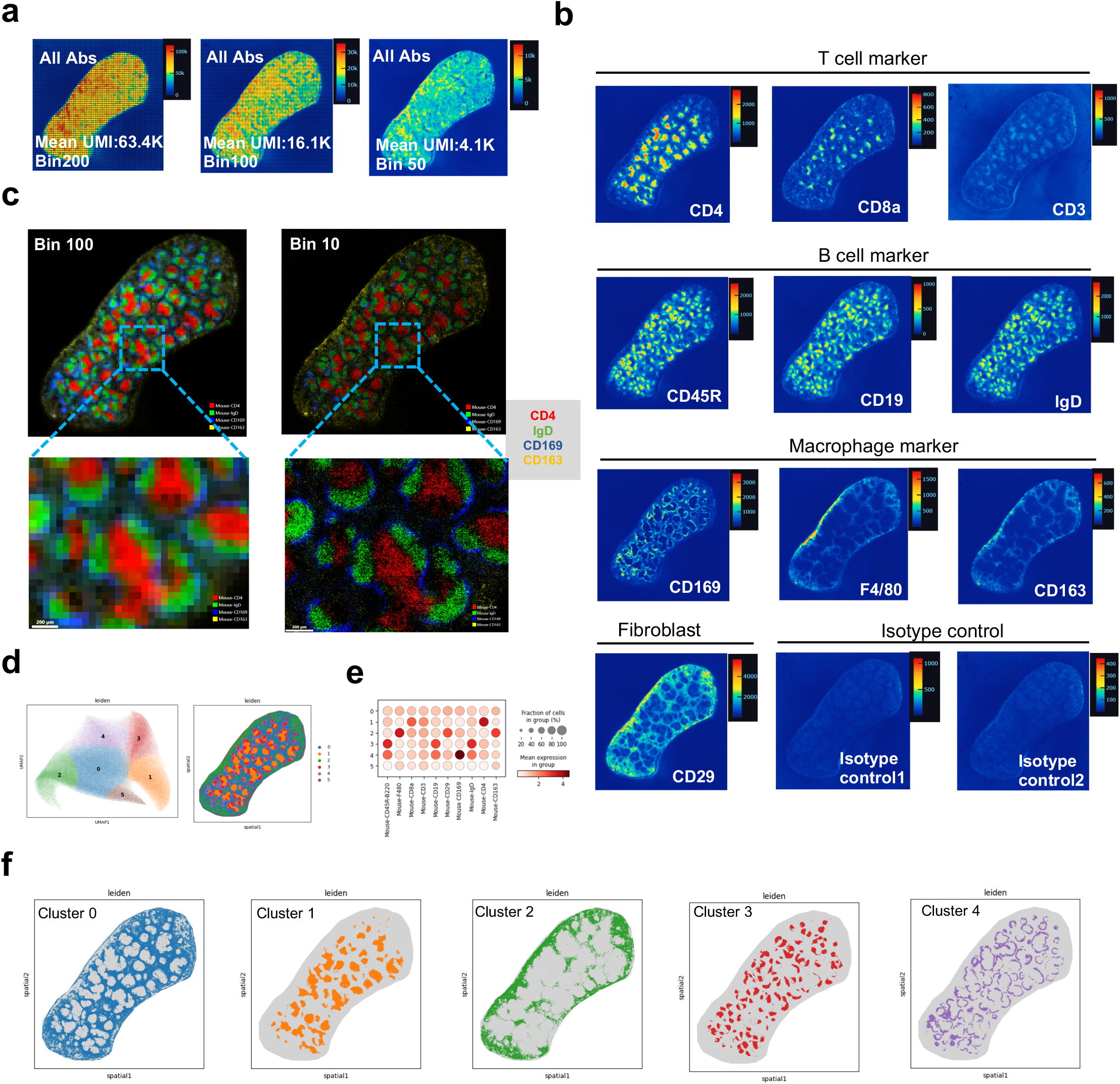
Stereo-CITE-seq analysis of mouse spleen. **a**, Spatial distribution of ADT UMI counts per Bin 200 (100 × 100 µm spatial area), Bin 100 (50 × 50 µm spatial area) and Bn 50 (25 × 25 µm spatial area). The average total ADT UMI counts per Bin 200, Bin 100 and Bin 50 were also shown. **b**, ADT expression patterns (UMI counts) of different surface markers of mouse spleen at Bin 10. **c**, CD4, IgD, CD169 and CD163 ADT expression patterns were assigned with different color and merged together to visualize their spatial location in the same image at Bin100 and Bin10 level. **d**, Non-supervised clustering of spatial barcodes of mouse spleen at Bin 10 level based on ADT expression. **e**, Dot plots showing ADT expression levels of identified clusters. **f**, Spatial location of identified clusters were shown.

Next, we performed Stereo-CITE-seq on mouse thymus tissues. We designed a 17-plex antibody panel to identify thymocytes (CD3, CD4, CD8a, CD27, CD5, CD44, CD90.2), myeloid cells (CD68, CD169, F4/80, CD11b, CD11c, Siglec-H), B cells (CD19, CD45R), fibroblasts (CD29) and endothelial cells (CD31). The mouse thymus can be broadly divided into the outer region known as cortex where immature thymocytes and stromal cells are densely packed, and the inner medulla region where the mature thymocytes and stromal cells are found at lower cell density ^28^. Our spatial ADT expression patterns faithfully recapitulated the expected thymus organization (Figure 3a). Thymus cortex region are characterized with higher expression of CD4+ CD8+ immature thymocytes as reported previously ^28^ (Figure 3a). CD5 is regarded as a negative regulator of the TCR signaling in immature thymocytes and a cell marker of medulla ^29,30^. We indeedobserved considerably high expression of CD5 in medulla region in our study (Figure 3a). CD44 is shown to be involved in invariant natural killer T cells (iNKT cells) development in medulla ^31^, which is confirmed in our study with higher expression levels in thymus medulla (Figure 3a). Macrophage subtypes were also identified through CD68 and CD169 ADT expression (Figure 3a). CD68-positive macrophages were found more abundant in cortex regions, while CD169-positive macrophages were more abundant within cortico-medullary junction (CMJ) (Figure 3a).

**Figure 3.**
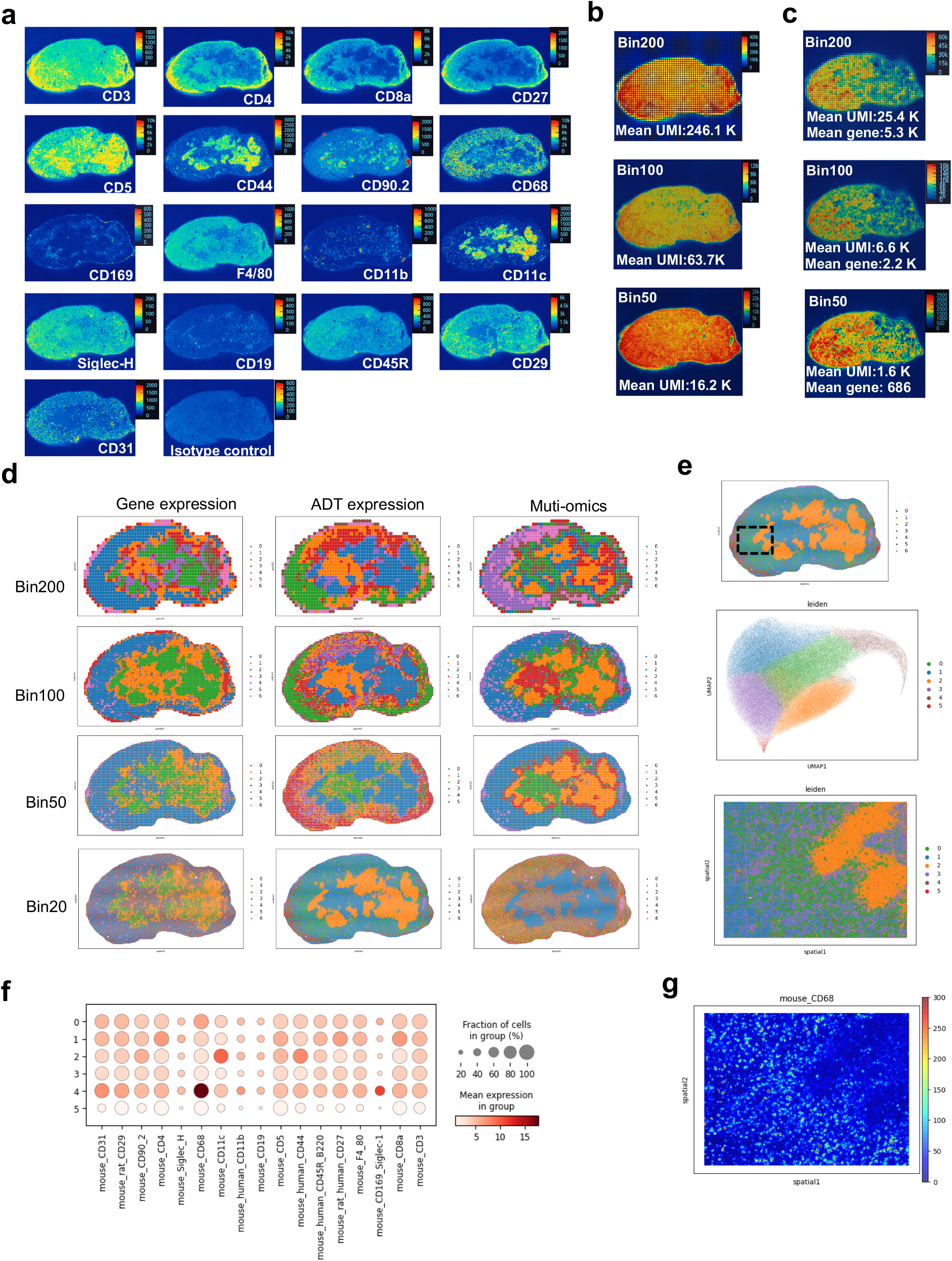
Stereo-CITE-seq analysis of mouse thymus. **a**, ADT expression patterns (UMI counts) of different surface markers for mouse thymus at Bin 50. **b**, Spatial distribution of ADT UMI counts per Bin 200 (100 × 100 µm spatial area), Bin 100 (50 × 50 µm spatial area) and Bn 50 (25 × 25 µm spatial area). The average total ADT UMI counts per Bin 200, Bin 100 and Bin 50 were also shown. **c**, Spatial distribution of gene expression (UMI counts) per Bin 200 (100 × 100 µm spatial area), Bin 100 (50 × 50 µm spatial area) and Bn 50 (25 × 25 µm spatial area). The average UMI counts and gene types per Bin 200, Bin 100 and Bin 50 were also shown. **d**, Non-supervised clustering of spatial barcodes based on gene expression, ADT expression and multi-omics data at different spatial resolution (Bin 200, Bin 100, Bin 50, Bin 10). **e**, Non-supervised clustering of spatial barcodes (Bin 5) of selected region of mouse thymus based on ADT expression. **f**, Dot plots showing ADT expression levels of identified clusters. **g**, CD68 ADT expression pattern was shown at selected regions of mouse thymus at Bin 5.

In the mouse thymus, we observed much higher total ADT UMI counts than mRNA UMI counts. The average protein UMI count per Bin 200 (100 × 100 µm area) is 246.1 K, whereas the average gene UMI count is 25.4 K, with almost 10-fold differences (Figure 3b and 3c). This could probably be due to the greater ease of capturing and reverse-transcribing ADT as compared to mRNA. The average protein UMI count per Bin 50 (25 × 25 µm spatial area) is 16.2 K (Figure 3b), while the average gene count per Bin 50 is 686 (Figure 3b), indicating that the workflow could be further optimized to increase mRNA capture efficiency. A previous study reported two macrophage populations within the thymus with distinct localization and origin ^32^. *Cx3cr1*+ macrophages are mainly located in medulla, while *Timd4*+ macrophages mostly reside in cortex ^32^. Consistent with that, *Cx3cr1*+ macrophages marker genes (*Ctsz, Cd63, Pmepa1* and *Zmynd15*) were found to be mainly expressed in thymic medulla region (Figure S1), while *Timd4*+ macrophage marker genes (*Hpgd, Serpinb6a, Slc40a1* and *Cd81*) were highly expressed in the cortex region (Figure S1).

Next, we performed non-supervised clustering of spatial barcodes based on gene expression, ADT expression and multi-omics data at various spatial binning resolutions (Figure 3d). Higher binning resolution such as Bin 50 and Bin20 could result in finer spatial distribution of clusters, while spatial distribution of resulting clusters at Bin 200 level could not faithfully recapitulate thymus structure (Figure 3d). Integrated analysis of ADT and gene expression using totalVI resulted in a much refined spatial cluster distribution especially at the CMJ region, at Bin50 in the Muti-omics group compared to Gene expression group (Figure 3d), substantiating the importance of protein information. We next focused on a smaller region in the thymus section, and performed non-supervised clustering at Bin 5 (Figure 3e). Among the identified 6 clusters, Cluster 4 is characterized with high level of CD68 expression, and cluster 0 expresses CD68 to a lower extent (Figure 3f). When we mapped these clusters to their spatial location, we discovered that Cluster 0 spatial barcodes were surrounding cluster 4 spatial barcodes (Figure 3e). Spatial pattern of CD68 ADT at Bin 5 resolution showed a clear separation of neighboring macrophages (Figure 3g). These results demonstrate that Stereo-CITE-seq could profile thymus residing macrophages at single cell resolution.

We next assessed the reproducibility and accuracy of Stereo-CITE-seq in mouse thymus, using the same 17-plex ADT panel as described above. We observed high correlation at bulk level between replicates for both the mRNA (Figure 4a) and protein expression (Figure 4b). We then performed non-supervised clustering of Bin 50 spatial barcodes based on gene expression (Figure 4c) and ADT expression (Figure 4e). We did not observe any significant difference in clustering distribution across replicates for both gene expression (Figure 4d and 4g) and ADT expression (Figure 4f and 4h), indicative of high reproducibility of the Stereo-CITE-seq platform. To evaluate the accuracy of Stereo-CITE-seq, we performed single marker immunofluorescence (IF) staining and proteomic profiling on the same tissue section (Figure 4i). IF intensity of markers of thymocytes (CD4, CD8 and CD44), endothelial cells (CD31), myeloid cells (CD11b and CD68) were highly corelated with respective the ADT expression patterns (Figure 4i), a testimonial to the robustness of Stereo-CITE-seq.

**Figure 4:**
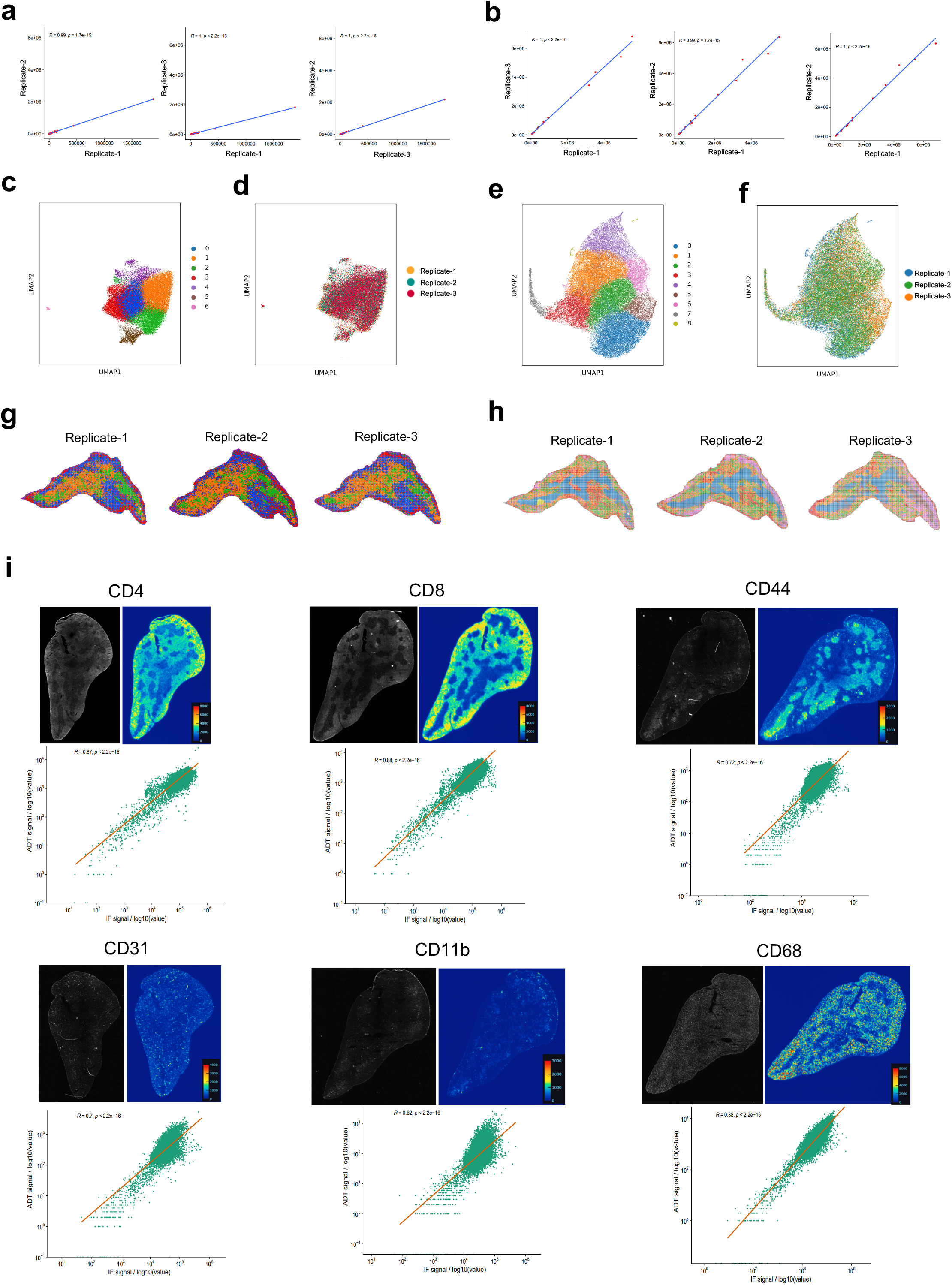
Reproducibility and accuracy of Stereo-CITE-seq in mouse thymus. **a**, Correlation analysis between all replicates of mRNA protein expression. Pearson correlation coefficient (R) and p-values were labeled on each panel. **b**, Correlation analysis between all replicates of 17 ADTs expression. Pearson correlation coefficient (R) and p-values were labeled on each panel. **c**, Non-supervised clustering of spatial barcodes (Bin 50) of all replicates based on mRNA expression. **d**, UMAP plot computed by gene expression, where colors indicate different replicates. **e**, Non-supervised clustering of spatial barcodes (Bin 50) of all replicates based on ADT expressions. **f**, UMAP plot computed by 17 ADTs expression, where colors indicate the different replicates. **g**, Visualization of spatial distribution of identified clusters in (c). **h**, Visualization of spatial distribution of identified clusters in (f). **i**, Correlation analysis between ADT UMI counts and IF intensities of the CD4, CD8, CD44 CD31, CD11b and CD68 across the spatial barcodes (Bin 50). Pearson correlation coefficient (R) and p-values were labeled on each panel.

Collectively, our data showed that we successfully developed a spatial multi-omic profiling method with high spatial resolution, reproducibility and accuracy, providing accurate co-detection of mRNAs and protein markers in biological samples.

## Methods

### Animal

All relevant procedures involving animal experiments presented in this study are compliant with ethical regulations regarding animal research and were conducted under the approval of the Animal Care and Use committee. Mouse thymus glands were dissected from 6-8 weeks old C57BL/6J male mice. After collection, tissues were embedded in Tissue-Tek OCT (Sakura, 4583) on dry ice and transferred to a -80°C freezer for storage before the experiment.

### List of antibodies used

**Table 1.**
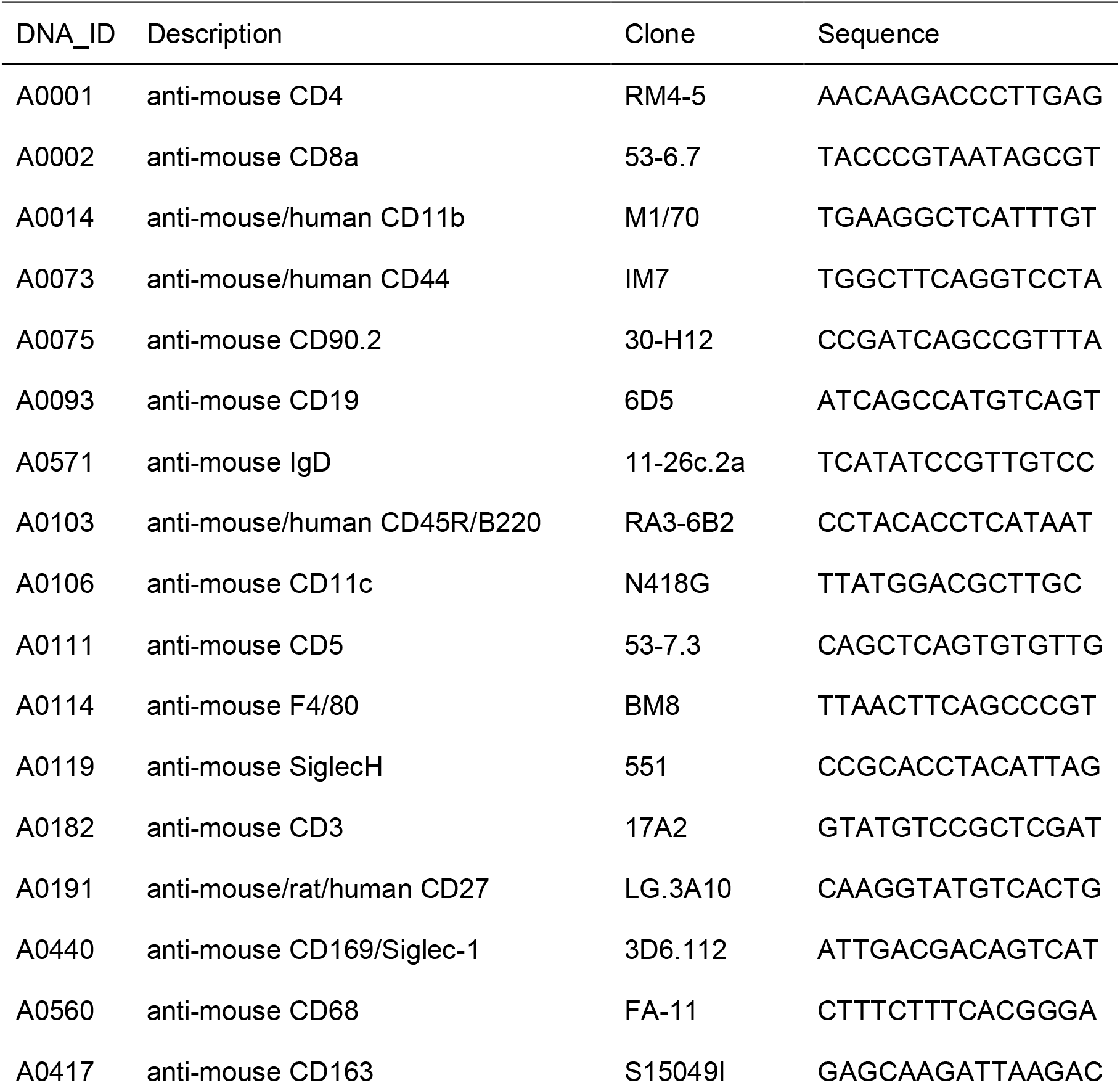
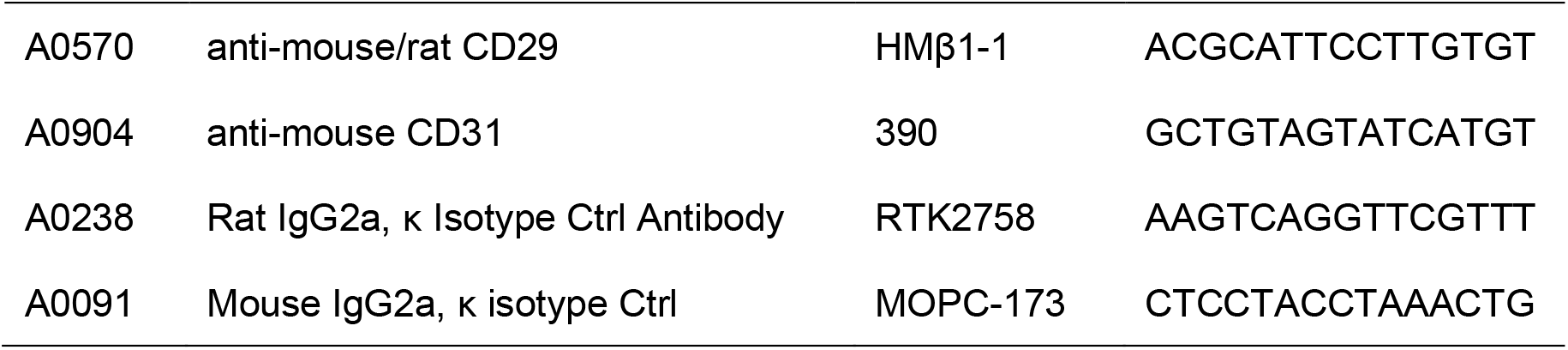
List of TotalSeq^™^ A anti-mouse antibodies used in this study

### Stereo-CITE-seq methodology

The following solutions were prepared freshly before experiment and were kept on ice. Blocking Buffer (50μl per sample): 3x SSC, 0.1% Triton-X-100, 10% serum, 1 μg/μl sheared salmon sperm, 2.5 μl TruStain FcX^™^ PLUS (anti-mouse CD16/32) Antibody (Biolegend, Cat. No.156604), 5% RNAse inhibitor (RI) and 5 μM 32nt Poly A. Wash Buffer: 3x SSC with 5% RI.

The OCT embedded tissue was cut in the cryostat at a thickness of the 10 μm and transferred onto a Stereo-seq chip. Next, the Stereo-seq chips with the tissue section were immediately warmed to 37°C for 1 minutes on a thermocycler adaptor. The tissue sections were fixed with pre-chilled methanol at -20°C for 30min. Next, the chips were blocked for 20 min with the Blocking Buffer and incubated with the TotalSeq™-A antibody mix in Blocking Buffer for 45 min at room temperature. The chips were washed three times with Wash Buffer for 30 seconds each. After DAPI staining, the chips were mounted with Glycerol and imaged. Next, sections were processed according to manufacturer’s instructions for Stereo-seq Transcriptomics T Kit with some modifications. In short, sections were permeabilized using Permeabilization Mix for an appropriate time determined by Stereo-seq Permeabilization Kit. After Reverse transcription, cDNA and antibody derived tag (ADT) products were released from the chips by Release Enzyme Mix and collected. The cDNA fraction was separated from ADT library by 0.8X beads selection, and was subjected to cDNA sequencing library preparation according to Stereo-seq Transcriptomics T Kit. The supernatant ADT product was cleaned up by 2.0X (0.8X+1.2X) beads selection and amplified using Protein PCR primer Mix. ADT and cDNA libraries were sequenced with MGI DNBSEQ-T7 or DNBSEQ-G400 Sequencer.

### Data processing

Gene expression and ADT Fastq files were generated by MGI DNBSEQ-T7 or DNBSEQ-G400 Sequencer. Stereo-seq transcriptomic raw data were processed as described previously ^9^. The pipeline was publicly available at https://github.com/BGIResearch/SAW. Stereo-seq ADT Fastq files were generated by MGI DNBSEQ-T7 or DNBSEQ-G400 Sequencer. CID and MID are contained in the read 1 (CID: 1-25 bp, MID: 26-35 bp) while the read 2 consist of 21 bp PCR handle and 15 bp antibody barcodes. CID sequences on the read 1 were first mapped to the designed coordinates of the *in situ* captured chip achieved from the sequencing, allowing 1 base mismatch. Reads with MID containing either N bases or more than 2 bases with quality score lower than 10 were filtered out. We used the in-house tag quantification pipeline to map antibody tags to their respective antibody barcodes. Finally, this information was used to generate a CID-containing ADT profile matrix.

### Transcriptomics and proteomics data integration

The expression profile matrix of mouse thymus was divided into bins of size n X n DNB, with n ∈ (20, 50, 100, 200). The transcripts of the same gene and the ADTs of the same protein were aggregated within each bin. Transcriptomic and proteomic expression matrices were clustered using SCANPY V1.9.1 ^33^. The transcriptomic data was preprocessed by adjusting gene expression, selecting highly variable genes, and performing PCA. Clustering was then performed using the leiden algorithm and visualized with UMAP. For the proteomic data, a nearest-neighbor graph was constructed, followed by leiden clustering and UMAP dimensionality reduction. Spatial visualization was performed with color-coded clusters. The joint analysis of transcriptome and proteome was performed using totalVI ^34^, and the resulting multi-omics model was clustered and visualized with UMAP and cluster spatial distribution.

### Evaluation of technical reproducibility and accuracy

Three consecutive thymus tissue sections were used to assess the reproducibility of the Stereo-CITE-seq. Gene and protein expression matric of three replicates were combined at bin size of 50 × 50 DNB for batch free clustering to compare the spatial distribution of clusters and across replicates. For the comparison of IF and ADT expression pattern. Secondary antibody was used to visualize the location of TotalseqA antibodies (CD4, CD8, CD44, CD31, CD11b and CD68) in each section. We normalized the ADT expression UMI count value and the IF staining intensity to 0-255 with spatial bins of 50 × 50 DNB. Then, Pearson’s correlation of the IF intensity and ADT expression on each slice were calculated.

## Data availability

The data that support the findings of this study have been deposited into CNGB Sequence Archive (CNSA)^35^ of China National GeneBank DataBase (CNGBdb)^36^ with accession number CNP0004285.

## Acknowledgements

The authors would like to acknowledge China National GeneBank for providing technical support.

## Competing Interests

A provisional patent related to this work has been filed by the BGI, and employees of BGI have stock holdings in BGI.

## Figure legends

**Figure S1.**
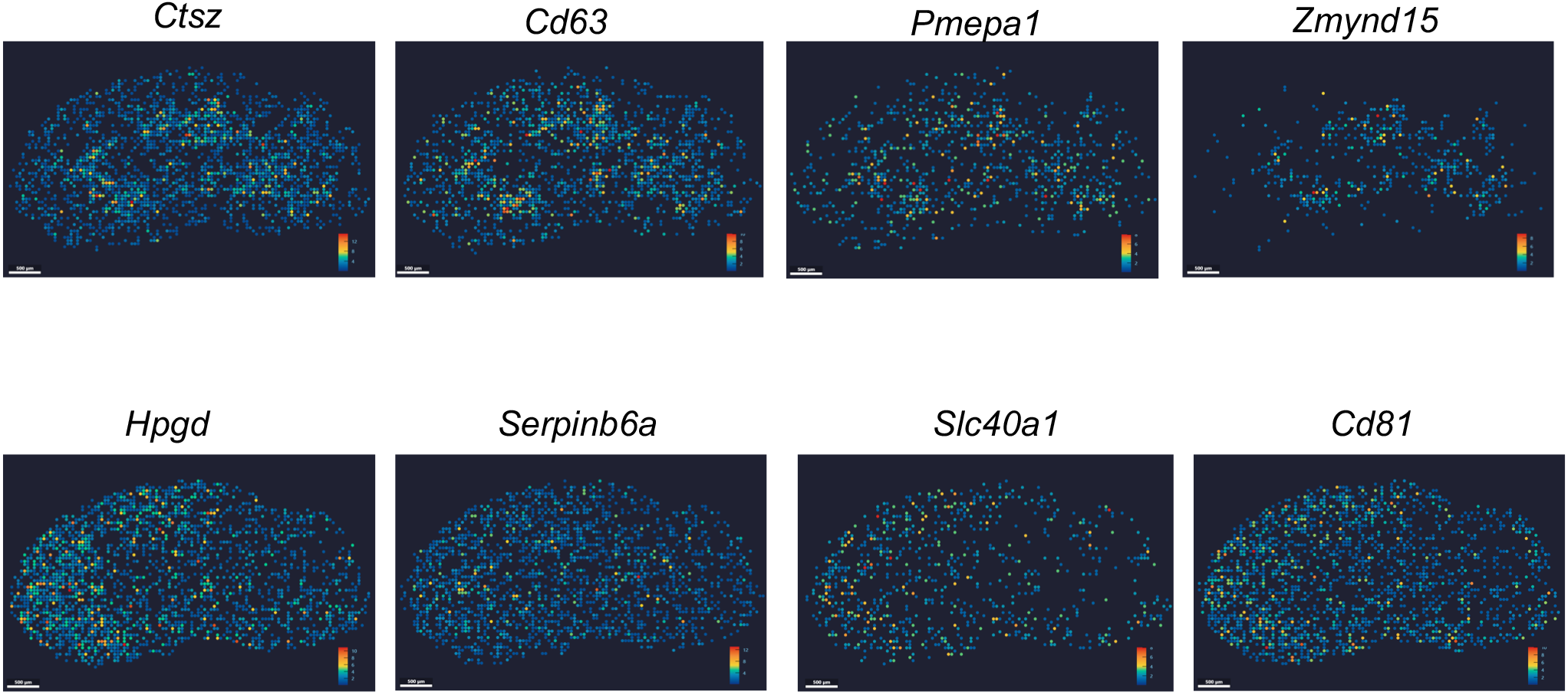
Stereo-CITE-seq confirms two macrophage subtypes with distinct spatial localization in the mouse thymus. Medulla-enriched *Cx3cr1*+ macrophage marker genes (*Ctsz, Cd63, Pmepa1, Zmynd15*) and medulla-enriched *Timd4*+ macrophage marker genes (*Hpgd, Serpinb6a, Slc40a1* and *Cd81*) were spatially visualized at Bin 100.

